# The Role of DOT1L Methyltransferase Activity in Fetal Hematopoiesis

**DOI:** 10.1101/2020.09.30.319889

**Authors:** Carrie A. Malcom, Joanna Piaseka-Srader, V. Praveen Chakravarthi, Shaon Borosha, Anamika Ratri, Nehemiah Alvarez, Jay L. Vivian, Timothy A. Fields, M.A. Karim Rumi, Patrick E. Fields

## Abstract

Early mammalian erythropoiesis requires the DOT1L methyltransferase. We demonstrated that loss of DOT1L in mutant mice resulted in lethal anemia during midgestation. The molecular mechanisms by which DOT1L regulates embryonic erythropoiesis have not yet been elucidated. In this study, a methyltransferase mutant mouse line (*Dot1L*-MM) was generated to determine whether the methyltransferase activity of DOT1L is essential for erythropoiesis. *Dot1L-*MM mice displayed embryonic lethality between embryonic days 10.5 and 13.5, similar to *Dot1lL* knockout (*Dot1L-*KO) mice. However, when examined at E10.5, unlike the *Dot1L-*KO, *Dot1L-*MM embryos did not exhibit evidence of anemia. In *ex vivo* hematopoietic differentiation cultures, *Dot1L-*KO and *Dot1L-*MM yolk sac (YS) cells both formed reduced numbers of myeloid, and mixed hematopoietic colonies. Erythroid colonies were able to be formed in numbers equal to wildtype embryos. Extensively self-renewing erythroblast (ESRE) cultures were established using YS cells from E10.5 embryos. *Dot1L-*KO and *Dot1L-*MM cells expanded significantly less than wild-type cells and exhibited increased cell death. Strikingly, *Dot1L-*KO and *Dot1L-*MM cells of YS origin exhibited profound genomic instability, implicating DOT1L methyltransferase activity in maintenance of the genome as well as viability of hematopoietic progenitors. Our results indicate that the methyltransferase activity of DOT1L plays an important role early murine hematopoiesis.

## 1. INTRODUCTION

Disruptor of telomere silencing 1-like (DOT1L) is essential for a number of biological processes during embryonic development, including early hematopoiesis (Feng et al., 2010). We observed that DOT1L deficiency in mice results in lethal anemia during mid-gestation (Feng et al., 2010). DOT1L mutant erythroid-myeloid progenitors failed to develop normally, showing a slow cell cycle progression associated with downregulation of GATA2, a transcription factor essential for erythropoiesis, and upregulation of PU.1, a factor that inhibits erythropoiesis (Feng et al., 2010). However, the precise molecular mechanisms underlying DOTL1L regulation of early hematopoiesis remain largely unknown.

Histone methylation is important for permissive or repressive chromatin conformation and can have a profound effect on the regulation of gene expression (Jambhekar et al., 2019). DOTL1L is the only known methyltransferase in eukaryotic cells responsible for the mono-, di- and tri-methyl marks on lysine 79 of histone H3 (H3K79) (Feng et al., 2002). These histone modifications are strongly associated with actively transcribed chromatin regions (Steger et al., 2008b). Despite the evolutionarily conserved nature of H3K79 methylation marks, the precise mechanism involved in H3K79 methylation-regulated gene expression remains unclear. DOT1L is a unique histone methyltransferase; unlike other histone lysine methyltransferases, the enzyme does not possess a SET domain (Min et al., 2003; Ng et al., 2002; Sawada et al., 2004; Singer et al., 1998). Instead, it contains a class I methyltransferase domain. Enzymes similar in structure are known to methylate DNA directly and can also target nonhistone proteins (Yang et al., 2013). The DOT1L substrate, H3K79 residue is also atypical in that it is localized on the nucleosome surface (Feng et al., 2002; Min et al., 2003; Sawada et al., 2004). These unique aspects of DOT1L-regulated H3K79 methylation and its consequences are incompletely understood and may impact the distinctive biologic role of the methyltransferase in early hematopoiesis.

In addition to early hematopoiesis, DOT1L is also involved in embryonic development of cardiac (Cattaneo et al., 2016; Pursani et al., 2018), and connective tissues (Djunaedi, 2018), as well as the regulation of a number of diverse cellular processes including, transcriptional elongation (Steger et al., 2008a), regulation of cell-cycle progression (Barry et al., 2009; Kim et al., 2014; Yang et al., 2019) and repair of DNA damage (FitzGerald et al., 2011; Kari et al., 2019; Wakeman et al., 2012). It remains unknown how H3K79 methylation is linked to cell differentiation required for the development of these specialized tissues. Moreover, the exact molecular details underlying DOT1L regulation of these cellular activities is unclear. DOT1L is quite a large protein (1540aa) that possesses multiple potential domains for interacting with many epigenetic and transcriptional regulators (Castelli et al., 2018; Chen et al., 2015a). However, the only molecular function that has been demonstrated for DOT1L is H3K79 methylation arising from the SAM-dependent methyltransferase (MT) fold (DOT1 domain) (Min et al., 2003; Ng et al., 2002; Sawada et al., 2004; Singer et al., 1998). Additional protein domains of DOT1L that can interact with other regulatory factors might also be linked to its diverse potential.

In this study, we sought to determine if DOT1L-mediated H3K79 methylation is dispensable for blood development. If this is the case, then the observed blood development defect in *Dot1L* knockout mice is a result of the loss of some function of the DOT1L protein that is distinct from its methyltransferase activity. Using CRISPR/Cas9, we generated a mouse line in which a point mutation resulting in a single amino acid substitution from asparagine to alanine was made in Exon 9 of *Dot1L* in mESCs (Asp241Ala). The mutation lies with the histone methyltransferase catalytic domain. Previous studies demonstrated that this substitution abolishes histone H3K79 methyltransferase activity (Min et al., 2003). Mutant mice contained an intact DOT1L protein that lacked H3K79 methyltransferase activity. The methyltransferase mutation was embryonic lethal-these mutant died around mid-gestation. The mice also displayed defects in embryonic hematopoiesis, including a decreased ability to form definitive myeloid, and multipotent (mixed) blood progenitors in *ex vivo* cultures. Unlike the *Dot1L* knockout (*Dot1L-* KO), yolk sac progenitors from the *Dot1L* methyltransferase mutant (*Dot1L-*MM) were able to produce erythroid colonies in numbers similar to wildtype. These results indicate that the histone methyltransferase activity of DOT1L plays a predominant role in the protein as a whole, at least in murine embryonic hematopoiesis.

## 2. MATERIALS AND METHODS

### 2.1. *Generation of Dot1L-*MM *and Dot1L-*KO *mutant embryonic stem cells*

Several mutant murine embryonic stem cells (mESC) were generated possessing mutations in the *Dot1* gene. To accomplish this, we utilized the CRISPR/Cas9 (Clustered Regularly Interspaced Short Palindromic Repeats/Cas9-associated) system (Chen et al., 2015b; Schwank et al., 2013). Using this system, along with single stranded DNA oligonucleotide repair templates, we seamlessly introduced two different point mutations into exon 9 of the *Dot1L* gene (schematic shown in **Figure 1**). The first of these mutations was a *Dot1L-*MM. This mutation changed a codon in exon 9, from AAT into GCT (shown in bold in **Figure 1C**). This change resulted in a single amino acid substitution from asparagine to alanine at amino acid 241 (Asn241Ala). Amino acid 241 lies within exon 9 in the methyltransferase domain of *Dot1L*. This asparagine is highly conserved and mutation resulting in this change has been shown to eliminate H3K79 methylation without changing DOT1L protein structure (Min et al., 2003). We also created 6 single nucleotide, silent mutations in the six codons immediately downstream, to enable clone screening via PCR. We screened for mESC in which both alleles were targeted with these mutations. A *Dot1L-*KO mESC was also generated which contained the same amino acid change and 6 silent mutations as in the *Dot1L*-MM. In addition, we added single nucleotide deletions at 22 and 53 nucleotides downstream of codon 241 in both alleles. These nucleotide deletions created two STOP codons, 12 and 19 amino acids downstream of the point mutation, which resulted in the generation of a non-functional, truncated DOT1L protein. The DNA sequences of the *Dot1L*-MM and *Dot1L*-KO mESC clones were verified by Sanger sequencing.

**Figure 1.**
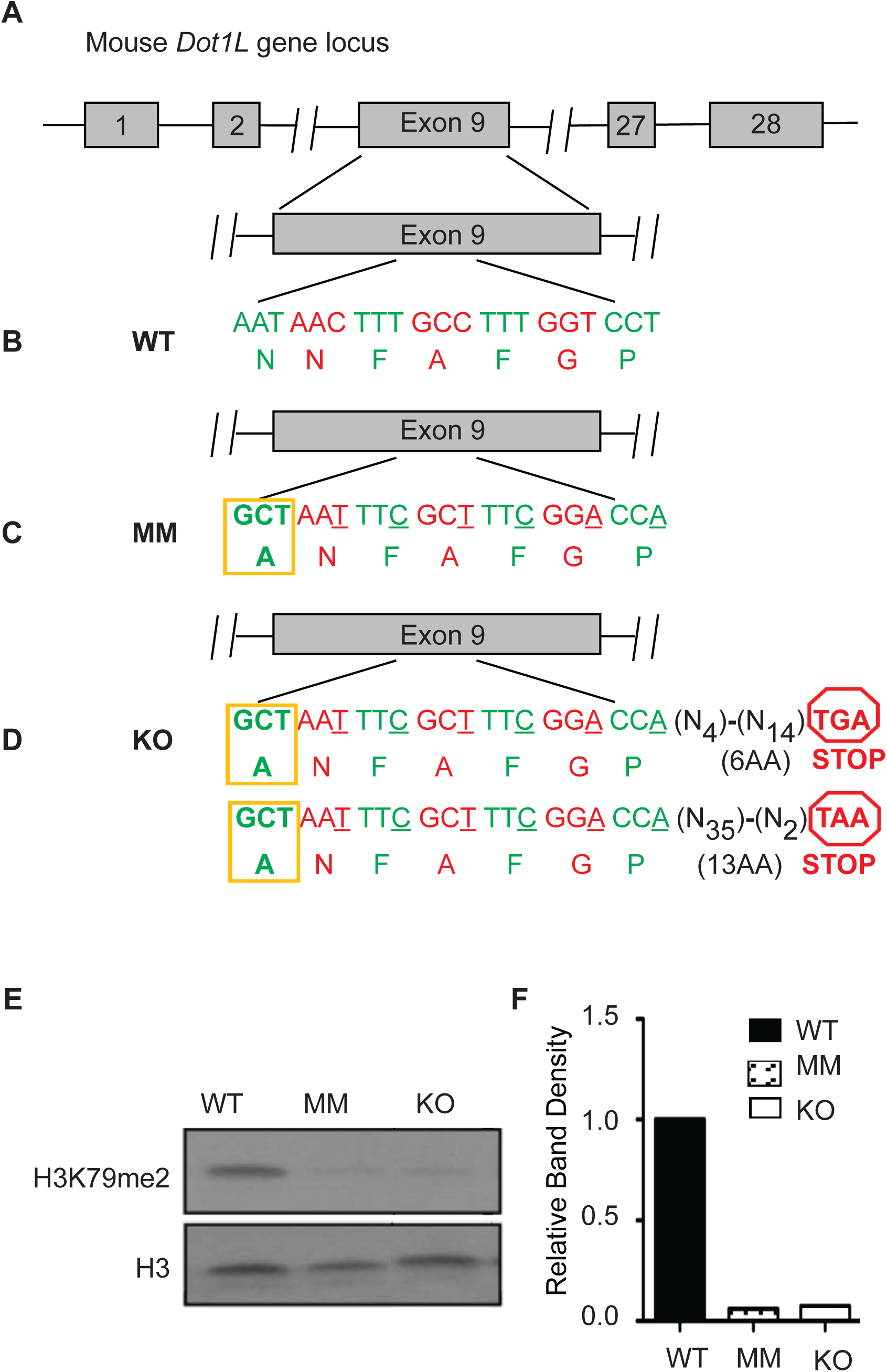
Generation of *Dot1L*-MM and *Dot1L*-KO embryonic stem cells. Murine ESCs were targeted with CRISPR/Cas9 to induce a point mutation Asp241Ala within Exon 9 of *Dot1L* gene (**A**-**C**). The MM clone contains this point mutation (in bold) in addition to 6 silent nucleotide changes (underlined) (**C**). The *Dot1L* knockout (KO) clone contains the point mutation and silent nucleotide changes as well as a single nucleotide deletion 53bp and 22bp downstream of the point mutation (**D**). These single nucleotide deletions create a stop codon shortly thereafter (**D**). Histones were extracted from wildtype (WT), MM, and KO mESC clones, and western blots were performed to detect H3K79 di-methylation (H3K79me2) (**E**). Histone H3 was detected as loading control (**E**). H3K79me2 band intensity is shown relative to that of WT mESC normalized with histone H3 band intensity (**F**).

### 2.2. Ex vivo *hematopoiesis with embryonic stem cell (mESC)-based culture system*

To differentiate mESCs into hematopoietic cells, mESCs were grown on tissue culture plates, then passage into a primary methylcellulose-based medium containing murine Stem Cell Factor (mSCF) following the manufacturer’s protocol (STEMCELL Technologies), where they differentiate into pluripotent, three dimensional structures called embryoid bodies (EBs) (Keller et al., 1993). EBs were then dissociated into a single cell suspension via a collagenase/trypsin mixture and re-plated into a secondary methylcellulose-based medium, containing cytokines that promote myeloid and erythroid differentiation [Interleukin (IL)-3, IL-6, and erythropoietin, (EPO)] (www.stemcell.com) (Keller et al., 1993). *Dot1L*-MM, *Dot1L*-KO, and wildtype mESCs were grown on MEF to prevent differentiation and then plated into the primary methylcellulose medium containing mSCF. mESCs were cultured at 3 different concentrations: 4000, 8000, and 12000 cells/35mm dish, where they were differentiated into embryoid bodies (EB) and counted on day 8.

### 2.3. Dot1L mutant mouse models

The *Dot1L*-KO mice were generated and maintained as described previously (Feng et al., 2010). To produce the *Dot1L*-MM mouse, we generated mutant mESC as described above. Among the mESC clones produced, we selected cells possessing the proper mutation in a single allele. The other allele was wildtype. Selected clones were given to the KUMC Transgenic and Gene Targeting Institutional Facility. They were expanded and karyotype normal cells were injected into blastocysts of C57/BL6 females to produce chimeras. *Dot1L-MM* heterozygous (*Dot1L*^+/MM^) F1 mice were generated by breeding chimeras with C57Bl/6 females (The Jackson Laboratory). Stocks of heterozygous mice were maintained by continuous backcrossing to C57Bl/6 stocks. Genotyping primers (described below) were used to identify mice possessing the mutated alleles(s) by PCR (**Figure 4B**). Genotyping was performed on ear punches that were digested overnight in a digestion buffer according to a protocol described previously (Wang and Storm, 2006). All animal experiments were performed in accordance with the protocols approved by the University of Kansas Medical Center Animal Care and Use Committee.

### 2.4. Mouse genotyping

Primers used for genotyping of the *Dot1L*-KO mice have been previously described (Feng et al., 2010). The following PCR primers were used to genotype *Dot1L*-MM mice:

*Methyl Mutation Fwd* (5’-GCTAATTTCGCTTTCGGACCA-3’)

*Methyl Mutation Rev* (5’-CTCCACAAGGGACAGCATGT-3’)

*Methyl Mutation Fwd WT* (5’-AATAACTTTGCCTTTGGTCCT-3’)

*Methyl Mutation Rev WT* (5’-CTCCACAAGGGACAGCATGT-3’)

The use of these primer pairs clearly identified the presence of *Dot1L*-MM allele and/or wildtype allele, respectively. Both PCR reactions produce bands ∼450bp in length.

### 2.5. Western blot analyses for detection of H3K79 methylation

Western blot analyses of H3K79 methylation (di-methyl H3K79) was performed on purified histones, whereas the expression of DOT1L was detected in total proteins extracted from mouse E10.5 embryonic fibroblasts (MEF). For histone extraction, MEFs were lysed in Triton Extraction Buffer [PBS with 0.5% Triton X-100, 2mM PMSF, 0.02% NaN3, and protease inhibitor cocktail (Roche)] at a density of 1×10^7^ cell/ml. Nuclear pellets were washed with the extraction buffer and resuspended in 0.2N HCl at a density of 4×10^7^ cell/ml, and histones were acid extracted overnight at 4°C. Cells lysates were centrifuged at 2000Xg for 10 minutes at 4°C, and supernatant containing histones was collected and stored at -80°C. Protein concentration was measured using the Bradford Assay. Proteins were separated on a 4-20% SDS-PAGE and transferred to PVDF membranes. An antibody against di-methyl H3K79 was obtained from Abcam, Inc (ab3594) whereas the other antibodies against unmodified histone H3, DOT1L or GAPDH were obtained from Cell Signaling Technologies, Inc. Binding of the primary antibodies were detected with HRP-conjugated, specific secondary antibodies. Signals of the immune complexes were detected by enhanced chemiluminescence reagents following standard protocols.

### 2.6. Ex vivo *erythroid-myeloid differentiation assays*

*Dot1L*-KO or *Dot1L*-MM heterozygous mutant males and females were set up for timed mating to collect the conceptuses on E10.5. Pregnant females were sacrificed, and uteri were dissected to separate embryos and yolk sacs (YS). Embryos were treated with RED extract-N-Amp Tissue PCR reagents (Millipore Sigma, Saint-Louis, MO) to purify genomic DNA and perform the genotyping PCR. YS were digested with collagenase I to dissociate the cells into single cell suspension and plated in methylcellulose medium (media M34334, StemCell Technologies) containing cytokines (SCF, IL-3, IL-6, and EPO) that promote definitive erythroid, myeloid, and mixed progenitor colony formation. The cells were cultured for 10 days, at which point colony number and area were assessed. Erythroid colonies included megakaryocyte/erythroid progenitors (MEP), blast forming unit-erythroid (BFU-E), and colony forming unit-erythroid (CFU-E). Myeloid colonies included bi-potential granulocyte/macrophage (CFU-GM) as well as singularly granulocyte and macrophage (CFU-G and CFU-M). Mixed colonies were multipotent, containing both erythroid and myeloid progenitors (CFU-GEMM).

### 2.7. Extensively Self-Renewing Erythroblasts (ESRE) assays

Digested E10.5 YSs were washed in IMDM alone, resuspended in 0.5ml expansion media for ESRE according to England et al., (England et al., 2011), and plated into gelatin-coated 24 well plates. Expansion media consisted of StemPro34 supplemented with nutrient supplement (Gibco/BRL), 2 U/ml human recombinant EPO (University of Kansas Hospital Pharmacy), 100 ng/ml SCF (PeproTech), 10^−6^ M dexamethasone (Sigma), 40 ng/ml insulin-like growth factor-1 (Pepro Tech) and penicillin-streptomycin (Invitrogen). After 1 day of culture, the nonadherent cells were aspirated, spun down, resuspended in fresh ESRE media, and transferred to a new gelatin coated well. After 3 days in culture, cells were counted on a hemocytometer using trypan blue exclusion. Day 3 after isolation is day 0 of expansion culture. Cells were then spun down, resuspended in fresh ESRE media, and plated into a new gelatin-coated well. Cells were counted every other day, resuspended in fresh media, and plated into 24 well plates. Cells were diluted via partial media changes to keep concentration at ≤ 2 x10^6^ cells/ml. Fold expansion was calculated relative to day 0 (day 3 after isolation).

### 2.8. Assessment of cell proliferation, cell cycle analyses and apoptosis assays

Single-cell suspensions from E10.5 YS were cultured in MethoCult™ GF M3434 (StemCell Technologies, Vancouver, BC, Canada) for 4 days. The mix of cytokines in this methylcellulose medium promotes definitive erythroid, myeloid, and mixed progenitor differentiation. Cells were collected on day 4 and stained with Annexin V to assess apoptosis. Some cells were fixed by adding cold 70% ethanol slowly to single cell suspensions, and then stained with propidium iodide (Belloc et al., 1994). Flow cytometry was performed by the use of a FACSCalibur (BD Biosciences, San Jose, CA)(Krishan, 1975). Analyses of the cytometric data were carried out using CellQuest Pro software (BD Biosciences) (Chiruvella et al., 2008; Sen et al., 2007; Sutherland et al., 2009).

### 2.9. Comet assays

An alkaline comet assay was performed using the Trevigen Comet Assay kit (Trevigen, Gaithersburg, MD). Cells were isolated from E10.5 YS of *Dot1L*-KO, *Dot1L*-MM and wildtype mice and grown in *ex vivo* erythroid differentiation media as described above. After 4 days of culture the cells were harvested by repeated pipetting resuspended in cold PBS. An aliquot of 1000 cells / 10µl was added to 100 µl of molten LMA agarose and spread onto a comet slide and an alkaline comet assay was performed following the manufacturer’s protocol. The slides were imaged using an upright Nikon Eclipse 80i fluorescence microscope at X20 and analyzed using CometScore Pro (TriTek Corp).

### 2.10. Statistical analyses

Each study group consisted of at least 6 individual mice. The experimental results were expressed as mean ± SE. Statistical comparisons between two means were determined with Student’s t-test while comparisons among multiple means were evaluated with ANOVA followed by Duncan *post hoc* test. *P* values ≤ 0.05 were considered as significant level of difference. All statistical calculations were done with SPSS 22 (IBM, Armonk, NY).

## 3 RESULTS

### 3.1 DOT1L methyltransferase activity is essential for ex vivo hematopoiesis from mESC

#### 3.1a. *Generation of Dot1L-*MM *and Dot1L-*KO *mutant embryonic stem cells*

Our previous work indicated that the Dot1L methyltransferase was necessary for early blood development. We wanted to determine whether the loss of the methyltransferase activity *per se* accounted for the blood development defect or if this observed reduction was due to some other previously unreported activity associated with the protein (either intrinsic or extrinsic). To first examine this, we generated several mutant murine embryonic stem cells (mESC) possessing mutations in the *Dot1L* gene, by utilizing the CRISPR/Cas9 system. Using this system, we seamlessly introduced two different point mutations into exon 9 of the *Dot1L* gene (schematic shown in **Figure 1A-D**). The first of these mutations was in the DOT1L methyltransferase domain (*Dot1L-*MM). This mutation changed a codon in exon 9, from AAT into GCT (**Figure 1B-C**) and resulted in a single amino acid substitution from asparagine to alanine at amino acid 241 (**Asn241Ala**). This asparagine is highly conserved and mutation resulting in this change has been shown to eliminate H3K79 methylation without changing DOT1L protein structure (Min et al., 2003). We screened for mESC in which both alleles were targeted. A *Dot1L-*KO mESC was also generated in which a STOP codon was introduced 12 amino acid downstream of the point mutation (Asn241Ala) in one allele, and 19 amino acid downstream in another, resulting in production of a non-functional, truncated DOT1L protein (**Figure 1D**).

#### 3.1b. Loss of H3K79 methylation in Dot1L-MM and Dot1l-KO mutant mESC

Histones were extracted from wildtype, *Dot1L*-MM, and *Dot1L-*KO mESC clones. Western blot analysis was performed to assess DOT1L methyltransferase activity by detection of H3K79 di-methylation. Both *Dot1L* mutants showed near absent H3K79 di-methylation relative to wildtype mESC (**Figure 1E**). Histone H3 was detected as loading controls. Comparison of H3K79 di-methylation band intensities demonstrated that the Asn241Ala point mutation ablated DOT1L histone methyltransferase activity in mESC similar to that of *Dot1L-*KO **(Figure 1E, F)**.

#### 3.1c. Dot1L Mutation has varying effects on embryoid body formation

Wildtype and *Dot1L*-MM mESC clones formed similar numbers of EB at all 3 concentrations of mESCs plated (**Figure 2A**). The *Dot1L*-KO clone formed fewer EB on average than wildtype and *Dot1L*-MM clones at all 3 concentrations; however, the differences were not statistically significant (**Figure 2A**). This growth deficiency only became apparent at the EB stage, as undifferentiated knockout mESCs had similar morphologies and growth rates as the other mESCs. Phenotypically, wildtype, *Dot1L*-MM, and *Dot1L*-KO EB resembled each other in size and shape (**Figure 2B-D**).

**Figure 2.**
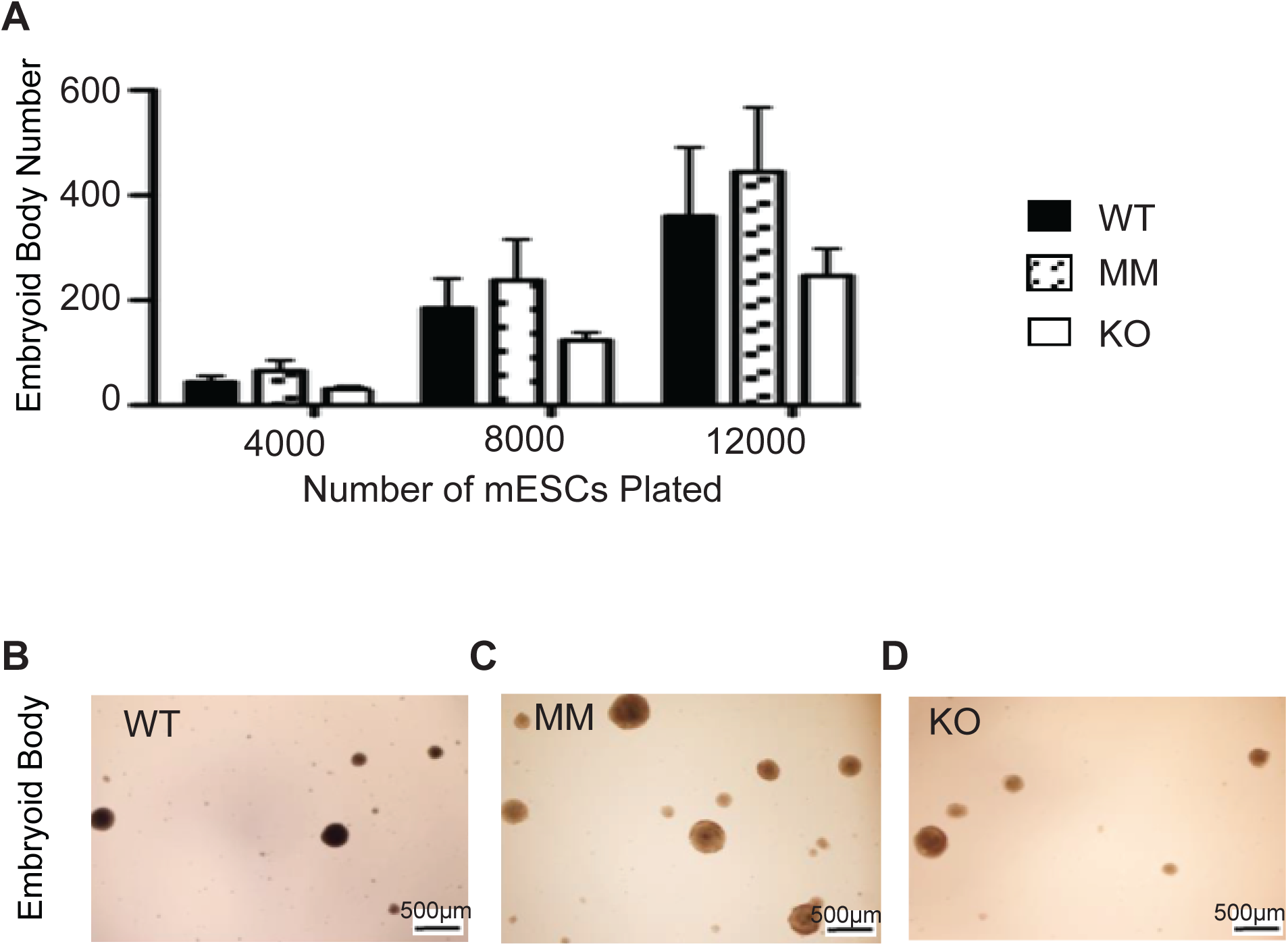
*Dot1L*-MM and *Dot1L*-KO had varying effects on embryoid body formation. Wildtype (WT), methyltransferase mutant (MM), and knockout (KO) mESC clones were cultured in methylcellulose medium containing mSCF for 8 days, where they differentiated to form embryoid bodies (EB). EB numbers were counted on day 8 (**A**). KO mESCs formed fewer EBs than WT and MM clones (**A**). However, the differences were not statistically significant. (**B-D**) Representative pictures of EBs formed from each mESC clone are shown.

#### 3.1d. *Loss of Dot1L methyltransferase activity impairs hematopoiesis* in vitro

Mutant, and wildtype EB were dissociated at day 8 into single cell suspensions and plated into secondary methylcellulose medium. This medium contained cytokines and growth factors that promote erythroid and myeloid hematopoietic differentiation. Day 8 EB cells have been shown to be ideal for formation of Blast Forming Unit-Erythroid (BFU-E, a definitive erythroid cell colony), myeloid colonies (both granulocyte and macrophage progenitors), and mixed colonies (multipotential colony containing both erythroid and myeloid progenitors) (Keller et al., 1993). They also have been shown to form immunophenotypic erythro-myeloid progenitors (EMP) and at peak levels along with definitive erythroid progenitors in some ES cell lines (McGrath et al., 2011). As EMP and definitive erythropoiesis emerge in the YS between E8.25-E11, day 8 embryoid bodies are good, *in vitro* representations of the E10.5 YS hematopoietic differentiation assays used to study *Dot1l* knockout mouse embryos (Feng et al., 2010). Primitive erythroid colonies (EryP) can also form from day 8 embryoid bodies, but they are not as predominant. Primitive erythropoiesis is better measured from day 6 embryoid bodies (McGrath et al., 2015). However, primitive erythroid colonies were noted if present in these cultures.

*Dot1L* knockout EB cells formed fewer primitive erythroid (EryP) and BFU-E colonies, on average, than wildtype (**Figure 3A**). This difference was not statistically significant. However, the knockout BFU-E colonies that did form were significantly smaller than wildtype (**Figure 3B**). Knockout EB cells also formed significantly fewer and smaller myeloid and mixed colonies than wildtype (**Figure 3A, C**). Likewise, *Dot1L-*MM EB cells formed fewer and smaller primitive erythroid and BFU-E colonies, on average, than wildtype (**Figure 3A, B**). *Dot1L-MM* EB cells also formed significantly fewer and smaller myeloid and mixed colonies compared to wildtype (**Figure 3A, C**). Interestingly, *Dot1L-*MM BFU-E colonies were larger on average than the knockout. Phenotypically, wildtype, *Dot1L-*MM EB, and *Dot1L-*KO EB cells formed BFU-E and myeloid colonies that were similar to wildtype in appearance, only smaller in size and with reduced cellularity (**Figure 3D-I**). Mixed colonies derived from mutant and knockout clones, however, were almost unrecognizable compared to wildtype (**Fig. 3J-L**). These colonies from both mutants contained both erythroid and myeloid cells, but at strikingly reduced numbers.

**Figure 3.**
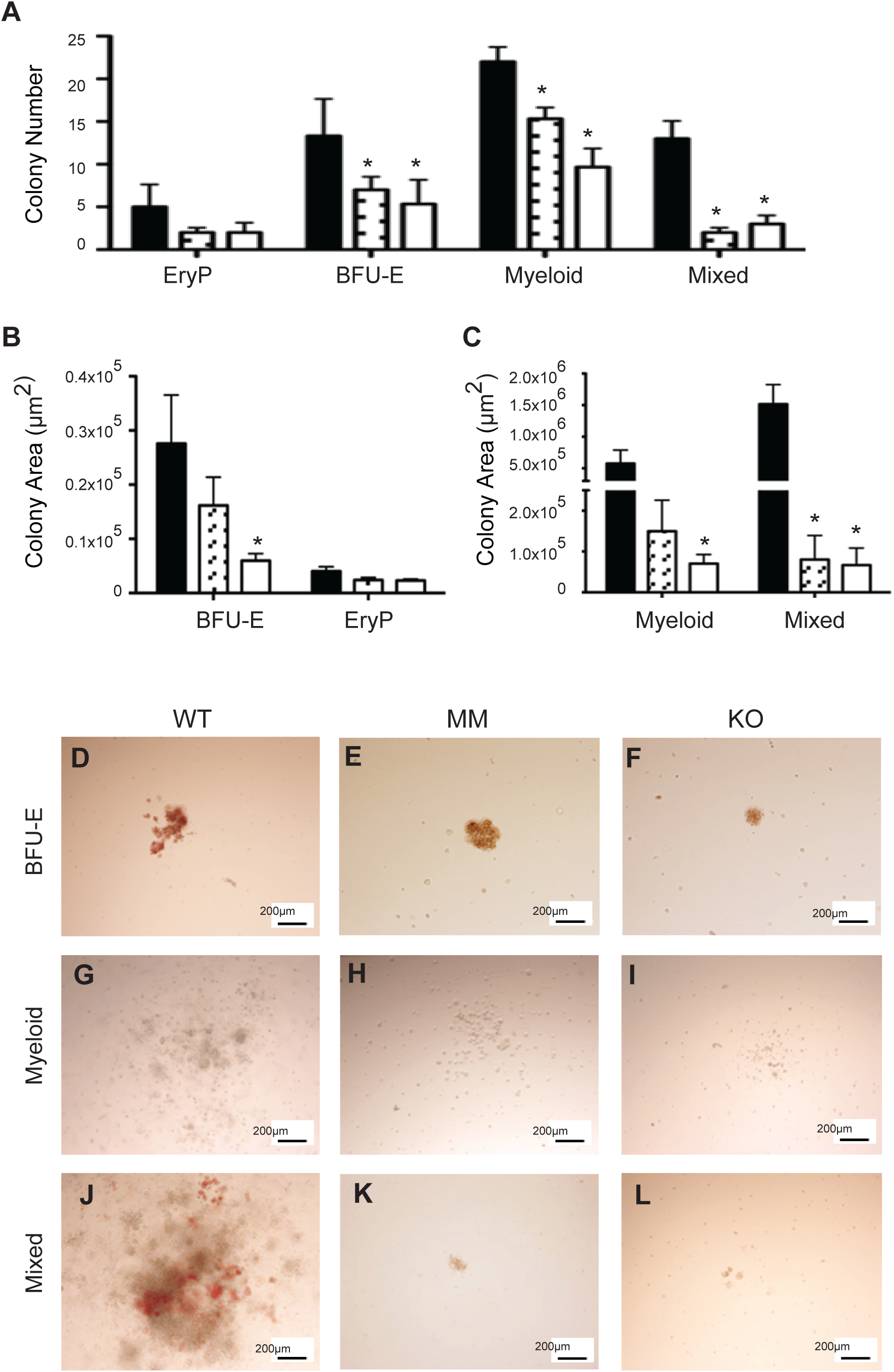
Loss of DOT1L methyltransferase activity impaired *in vitro* hematopoiesis. Equal numbers of wildtype (WT), methyltransferase mutant (MM) and knockout (KO) day 8 embryoid body (EB) cells were cultured in methylcellulose medium containing cytokines that promote primitive erythroid (EryP), definitive erythroid (BFU-E), myeloid (granulocyte, macrophage), and mixed (multipotential, erythroid and myeloid) differentiation. EryP, BFU-E, myeloid, and mixed colonies were counted (**A**) and area measured (**B** and **C**) on day 10. *Dot1L*-MM and *Dot1L*-KO mESCs, on average, formed fewer and smaller hematopoietic colonies compared to WT. Representative pictures of BFU-E (**D-F**), myeloid (**G-I**), and mixed (**J-L**) colonies are shown in the lower panels of the figure.

### 3.2. Development of the Dot1L-MM mouse: the methyltransferase activity of DOT1L is required for optimal hematopoiesis

The generation and characterization of the *Dot1L-KO* mutant mice has been described previously (Feng et al., 2010). *Dot1l-KO* mice express very low levels of *Dot1L* mRNA and the truncated DOT1L protein is not detected (Feng *et al.*, 2010). Loss of DOT1L expression in *Dot1L-* KO mice resulted in complete loss of the mono, di- and tri-methyl marks on H3K79 (Feng *et al.*, 2010). Dot1L-MM were generated by introducing a point mutation (Asn241Ala) in exon 9 that encodes part of catalytic domain (**Figure 4A**). The point mutation was detected by an allele specific PCR genotyping (**Figure 4A, B**). Although *Dot1L-*MM mice expressed a normal level of DOT1L protein (**Figure 4C, D**), the DOT1L-MM protein lacked the methyltransferase activity required for H3K79 methylation (**Figure 4E, F**). For the current analysis, we compared *Dot1L*-KO mice and *Dot1L*-MM mice each with wildtype control mice from the same litter.

**Figure 4.**
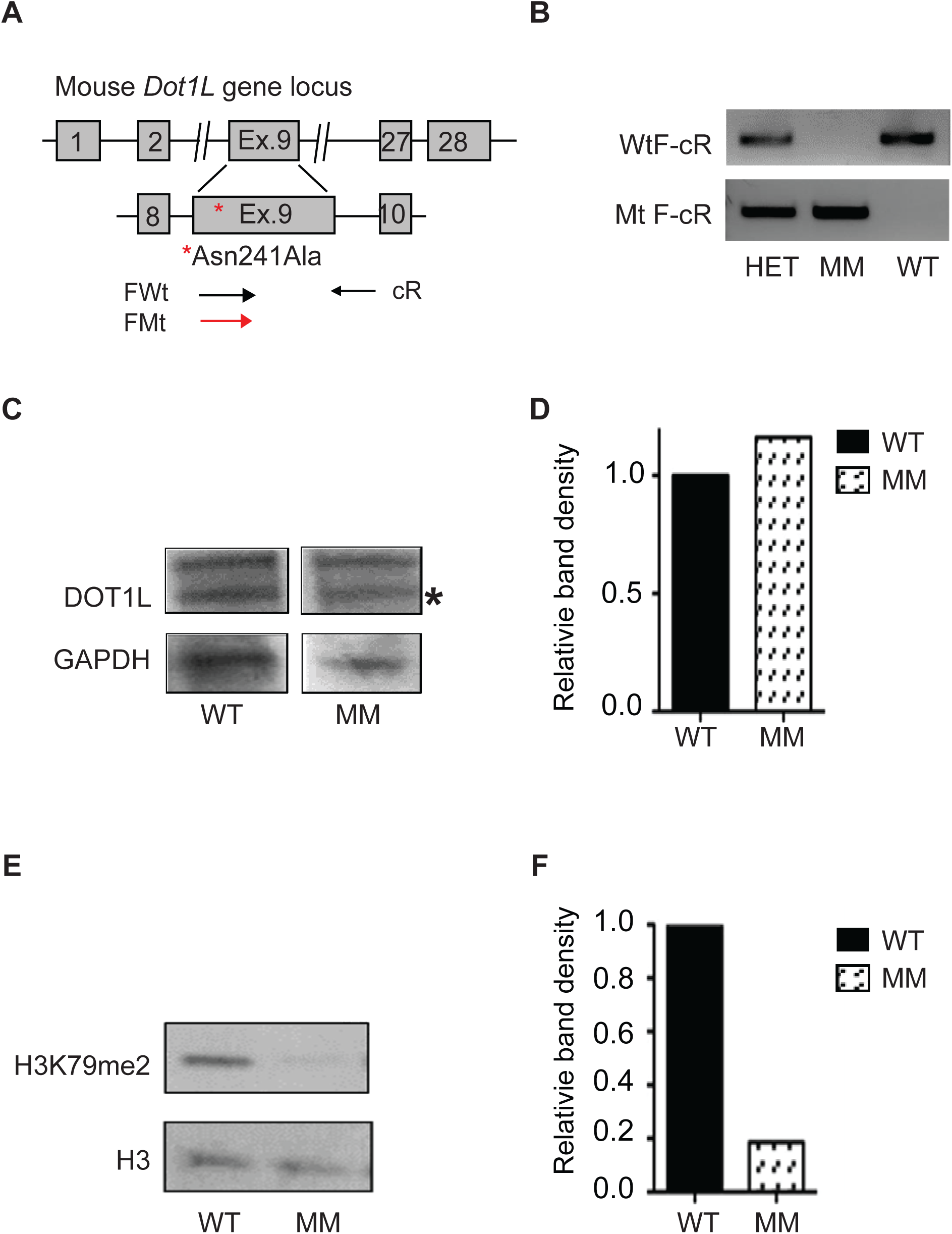
Generation of *Dot1L*-MM mice. (**A**) Schematic diagram showing the region of mouse *Dot1L* gene targeted by CRISPR/Cas9 to induce the point mutation Asn241Ala that eliminates H3K79 methyltransferase activity and the allele specific primers used to screen for mutants. (**B**) PCR genotyping using the primers shown in (**A**) to identify wildtype (WT), heterozygous (Het), and methyltransferase mutant (MM) embryos. Histones and whole protein lysates were extracted from E10.5 MEFs and western blot analyses were performed for histone H3, histone H3 lysine 79 di-methylation (H3K79me2) and DOT1L, GAPDH, respectively (**C-F**). Band densities were normalized to loading controls (histone H3 and GAPDH) and are shown as relative to wildtype (**D, F**).

#### 3.2a. Embryonic lethality of Dot1l mutant mice

Heterozygous *Dot1L-KO* or *Dot1L-MM* males and females were bred together, and embryos were collected and genotyped (described in Material and Methods) on E10.5, E11.5, E12.5, E13.5 and postnatal day 21 (PND21) to assess their viability. Viability of knockout embryos has been shown in our previous report (Feng et al., 2010). At E10.5 *Dot1L-*MM embryos showed normal Mendelian ratios. About 10% of MM embryos were dead at E11.5. None of the *Dot1L-* MM embryos were alive at E13.5 or PND21 (**Figure 5A**). Prior to their death, *Dot1L-*KO embryos were smaller in size, displayed abnormalities in heart development, altered vasculature, and a severe anemia (Feng et al., 2010). *Dot1L-*MM embryos at E10.5 were slightly smaller than their wildtype littermates but larger than *Dot1L-*KO embryos (**Figure 5B-D**). *Dot1L-*KO embryos were much smaller and severely anemic as previously reported (Feng et al., 2010) (**Figure 5D, G)** but *Dot1L-*MM YS were well vascularized and contained blood (**Figure 5C**). Remarkably, the *Dot1L-* MM embryos contained almost normal amounts blood, which was visible in the heart and aorta-gonad-mesonephros (AGM) region (**Figure 5C** and **F**).

**Figure 5.**
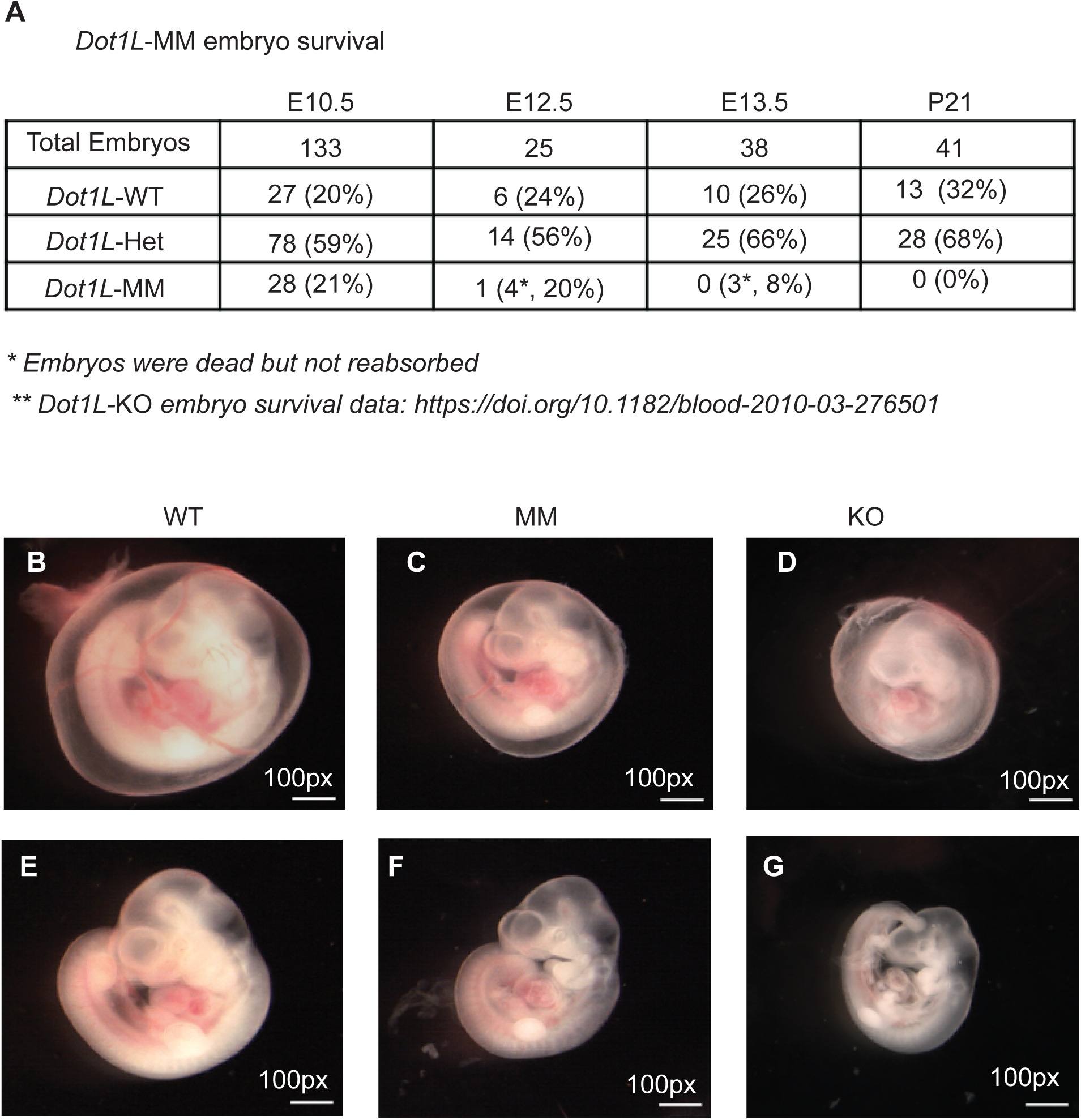
Embryonic lethality of *Dot1L*-MM mice. *Dot1L* methyltransferase mutant (*Dot1L*- MM) embryos display embryonic lethality but minimal defects in growth and hematopoiesis. (**A**) Table showing the percentage of wildtype (WT), *Dot1L*-MM heterozygous (Het), and homozygous methyltransferase mutant (MM) embryos alive at embryonic days 10.5, 12.5, and 13.5, and 21 days after birth (P21). *Dot1L*-MM embryos die between E10.5 and E13.5 (**A**). Representative pictures of WT, *Dot1L*-MM, and *Dot1L*-KO E10.5 embryos are shown in the image panels (**B-G)**. *Dot1L*-MM YSs have slightly less prominent blood vessels than WT but higher than that of *Dot1L*- KO (**B-D**). *Dot1L*-MM embryos are also slightly smaller and mildly anemic compared to WT embryos, but they are not as small or anemic as *Dot1L*-KO embryos (**E-G**).

#### 3.2b. DOT1L methyltransferase activity is essential for optimal ex vivo hematopoiesis of YS cells

E10.5 YS and their corresponding embryo tissue were obtained from *Dot1-L* MM mice by timed mating between *Dot1L-*MM heterozygous mice (*Dot1L-*^*+/M*^ X *Dot1L-*^*+/M*^). Likewise, similar timed mating was carried out to obtain *Dot1L-*KO tissues. DNA was extracted from corresponding embryo tissue and used for genotyping. YS were digested in 0.5ml digestion buffer and counted. Comparable numbers of YS cells from all mouse lineages were plated. Knockout YS cells formed fewer erythroid colonies than wildtype, while MM cells were able to form similar numbers. Production of myeloid lineage colonies from *Dot1L-*MM YS cells was reduced by approximately 50%, while those formed from knockout cells was reduced by approximately 75% (**Figure 6A)**. Strikingly, from both mutants there was approximately a 90% reduction in the formation mixed colonies, when compared to cells from wildtype littermates (**Figure 6A**). These colonies contain multipotential cells that can differentiate into either erythroid or myeloid progenitors. The areas of the colonies were calculated. Erythroid colony area was significantly less in colonies formed from cells from *Dot1L-*MM YS, and further diminished in those formed from *Dot1L-*KO cells (**Figure 6B, C**). The average size of the myeloid colonies formed by cells from both mutants was similar (**Figure 6B, D**). Those cells from mutant YSs that did form mixed colonies produced colonies of similar average size (**Figure 6B, E**).

**Figure 6.**
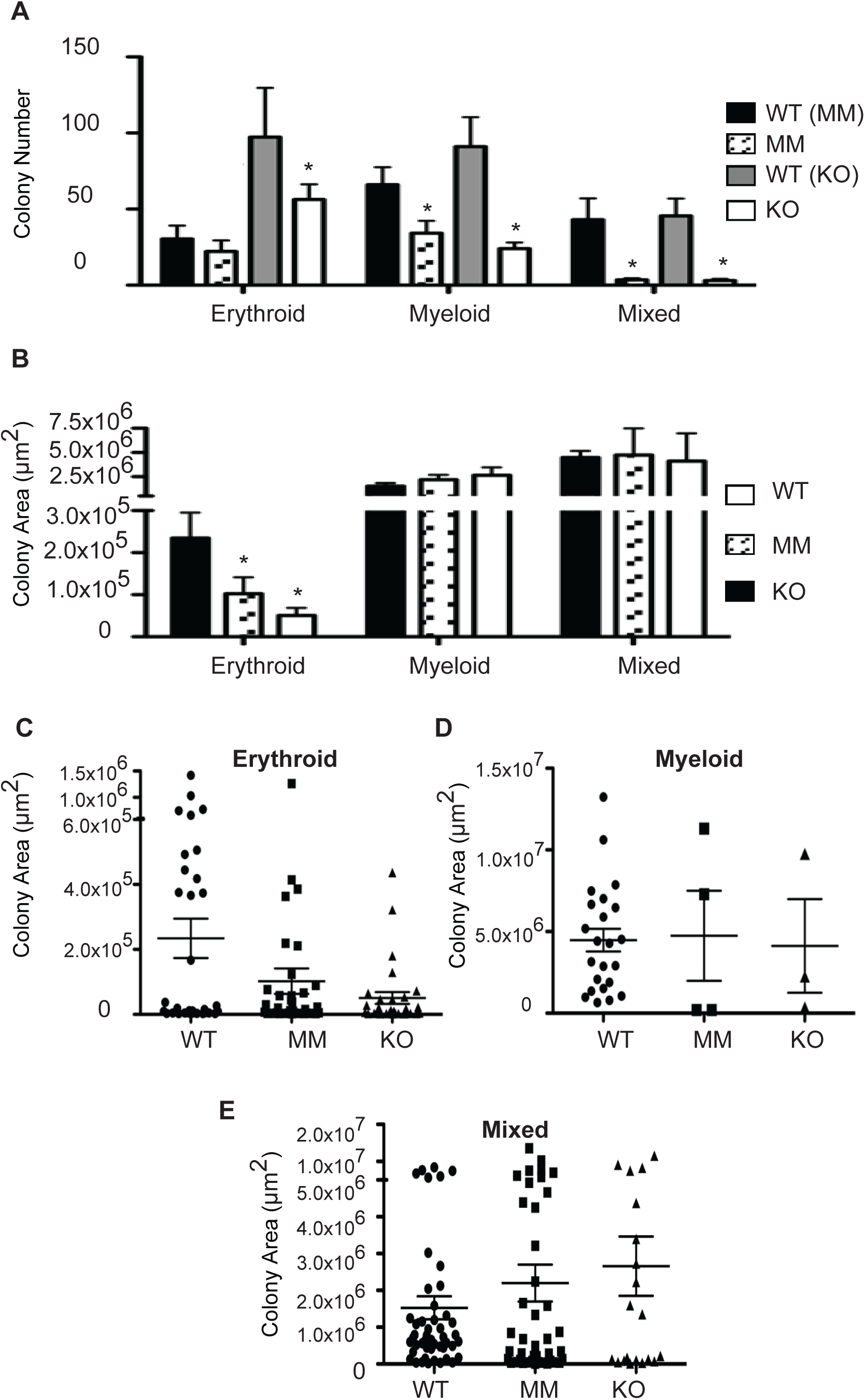
*Dot1L*-MM progenitors from YS displayed altered hematopoiesis. Equal numbers of wildtype (WT) and methyltransferase mutant (MM), wildtype from *Dot1L* knockout (KO) E10.5 YS cells were cultured in methylcellulose medium containing cytokines that promote definitive erythroid (megakaryocyte/erythroid, BFU-E, and CFU-E progenitors), myeloid (granulocyte/macrophage progenitors), and mixed (multipotential, erythroid and myeloid progenitors) differentiation. Erythroid, myeloid, and mixed colonies were counted (**A**) and area measured (**B**) on day 10. *Dot1L*-MM YS formed similar numbers of erythroid colonies as WT, but they were significantly smaller in size (**B**). Cells from *Dot1L*-MM YS formed significantly fewer myeloid and mixed colonies than that of WT, but the few colonies that did form achieved similar sizes as WT (**B**). Dot plots showing the distribution of erythroid (**C**), myeloid (**D**), and mixed (**E**) colony areas of E10.5 WT, *Dot1L*-MM, and *Dot1L*-KO YS cells after the hematopoietic differentiation assays.

When colonies that were formed by cells from either *Dot1L-*MM or *Dot1L-*KO mutant YS were examined by light microscopy, there was an obvious feature that they all had in common. Although the average areas and distribution of areas of the colonies were similar, the mutants had significantly less cellularity (far fewer cells per colony) (**Figure 7D-F** and **J-L)** than the corresponding colonies formed by wildtype YSs (**Figure A-C** and **G-I)**. This is clearly evident in the images of representative colonies shown in this figure (**Figure 7A-L**).

**Figure 7.**
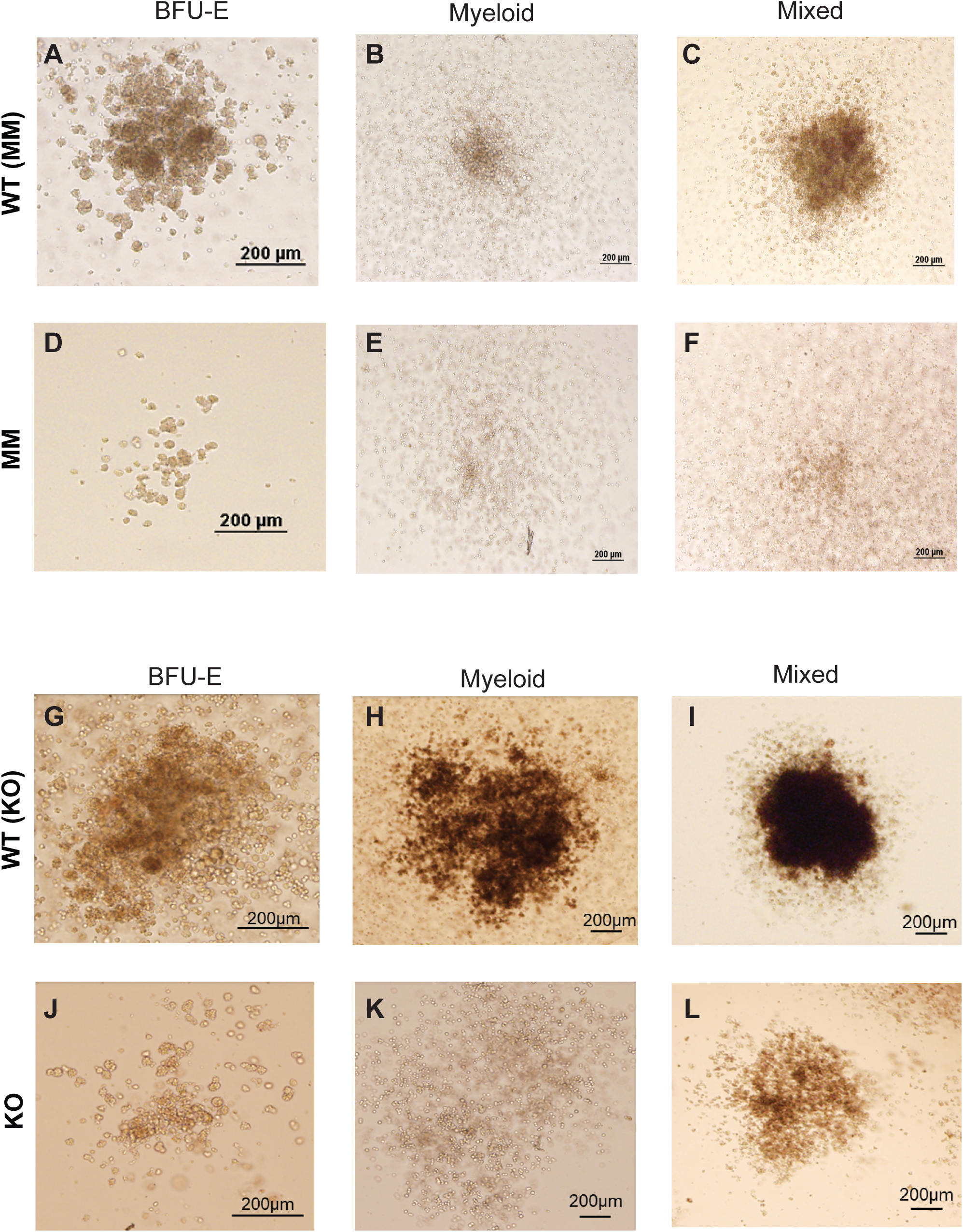
Erythroid, myeloid, mixed colonies from YS progenitor cell culture. Wildtype (WT), methyltransferase mutant (MM), *Dot1L* knockout (KO) E10.5 YS cells were cultured in methylcellulose medium containing cytokines that promote definitive erythroid (megakaryocyte/erythroid, BFU-E, and CFU-E progenitors), myeloid (granulocyte/macrophage progenitors), and mixed (multipotential, erythroid and myeloid progenitors) differentiation. Representative pictures of *Dot1L*-MM BFU-E (**A** and **D**), myeloid (**B** and **E**), and mixed (**C** and **F**) colonies are shown in the upper panels. In the lower panels, representative pictures of *Dot1L*- KO BFU-E (**G** and **J**), myeloid (**H** and **K**), and mixed (**I** and **L**) colonies are shown.

#### 3.2c. Dot1L mutant erythroblasts displayed decreased cell proliferation, defective cell cycle progression and increased apoptosis

Cells isolated from mutant and wildtype YSs were cultured in expansion medium and differentiated into ESRE. ESRE cells were counted every other day for 2 weeks to calculate cell proliferation. Both *Dot1L*-MM and *Dot1L*-KO erythroblasts had severely blunted proliferation, which resulted in a significantly reduced number of growing cells compared to wildtype (**Figure 8A, B**). Cell cycle analysis revealed that a larger proportion of both *Dot1L*-MM and *Dot1L*-KO progenitors displayed a G_0_/G_1_ cell cycle arrest (**Figure 8C-G**). A greater percentage of *Dot1L*- MM and *Dot1L*-KO YS hematopoietic progenitors were undergoing apoptosis, compared to wildtype, as measured by Annexin V positivity (**Figure 8H**). But the percentage of apoptosis was even higher in *Dot1L*-KO compared to *Dot1L*-MM YS derived cells (**Figure 8H).** These data suggest that the mutant YS blood progenitors are not proliferating or surviving in hematopoietic differentiation cultures to the same extent as wildtype.

**Figure 8.**
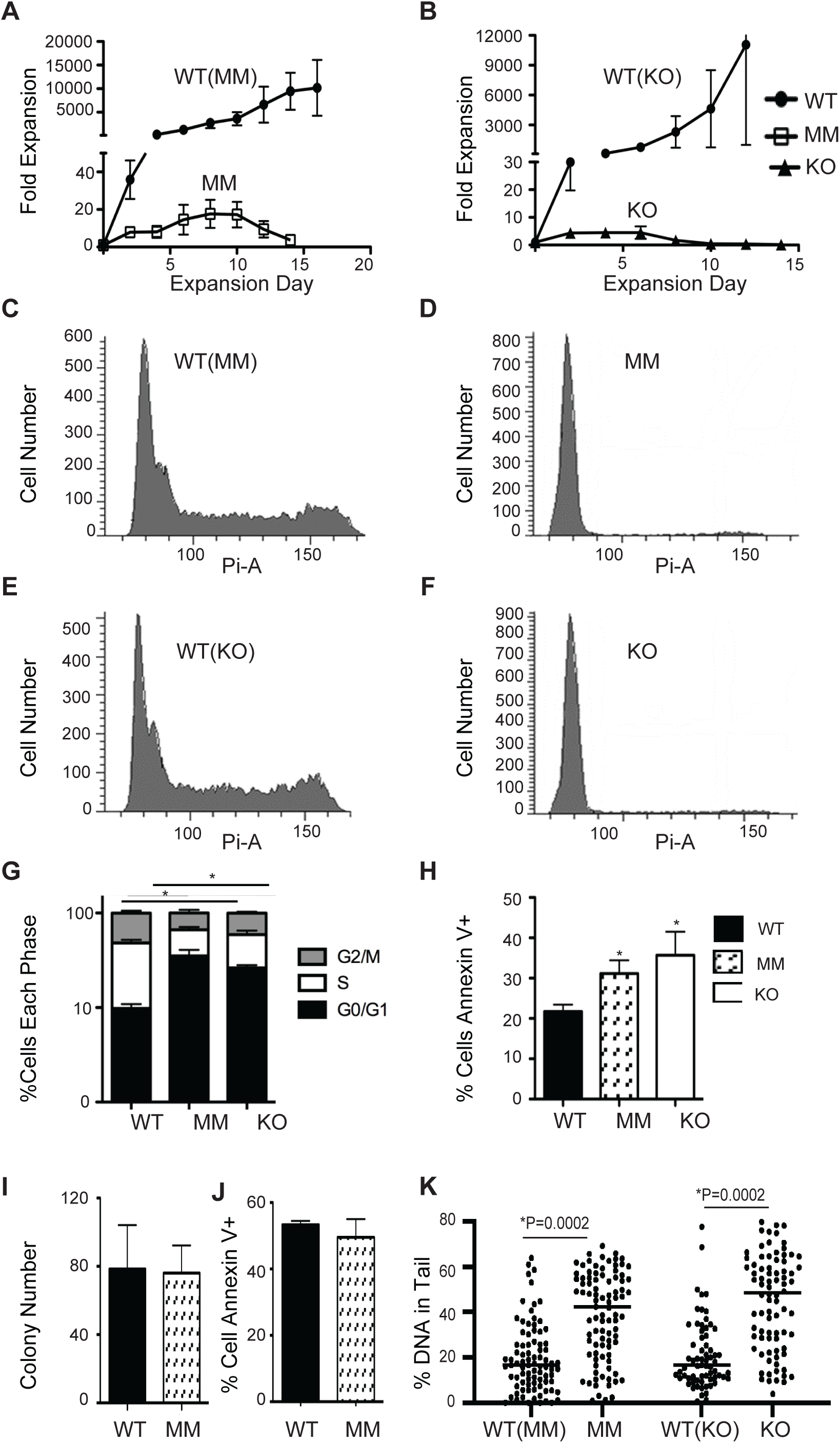
*Dot1L*-MM and *Dot1L*-KO cells exhibited decreased proliferation, cell cycle arrest, increased apoptosis and higher DNA damage. Extensively self-renewing erythroblast (ESRE) cultures were performed using E10.5 YSs from *Dot1L*-MM (**A**) and *Dot1L-KO* (**B**). The data showed that cells from *Dot1L*-MM and *Dot1L-KO* YSs display almost similar phenotypes regarding proliferation (**A, B**). In the ESRE cultures, *Dot1L*-MM and *Dot1L*-KO YS cells formed significantly fewer ESREs and had increased cell death compared to wildtype (**A, B**). E10.5 YS cells were cultured in methylcellulose medium containing cytokines that promote definitive erythroid, myeloid, and mixed progenitor differentiation. Cells were collected on day 4 and stained with propidium iodide to asses cell cycle progression in WT-MM (**C**), *Dot1L*-MM (**D**), WT-KO (**E**) and *Dot1L*-KO (**F**). Percentage of cells in G2/M, S, G0/G1 phase in WT, MM and KO showed that most of the cell in MM and KO were accumulated in G0/G1 phase (**G**). Cells were collected on day 4 were also stained with Annexin V to assess apoptosis. Compared to wildtype, a greater percentage of hematopoietic progenitors from *Dot1L*-MM and *Dot1L*-KO YS were found to be apoptotic, as measured by Annexin V positivity (**H**). To determine the role of DOT1L methyltransferase activity, *Dot1l-*MM E10.5 YS cells were plated in methylcellulose medium containing cytokines that promote *only* mature erythroid colony formation (mature BFU-E and CFU-E). Interestingly, no differences were found in colony number between *Dot1L-*MM and wildtype cells (**I**). After 4 days in culture, mature erythroid cells were labeled with Annexin V and analyzed via flow cytometry for apoptosis. No differences were found in the percentage of cells undergoing apoptosis between *Dot1L-*MM and WT cells. (**J**). In addition, Alkaline Comet assays were performed on day 6-8 ESREs to assess DNA damage; *Dot1L* MM and KO ESREs showed greater DNA damage than WT (**K**).

The above-mentioned experiments were designed to determine whether DOT1L activity was necessary for growth and survival of all myeloid and erythroid lineages. Thus, the YS cells were cultured in conditions (with specific cytokines) that enabled the growth of both erythroid, myeloid and mixed lineages. We previously reported that the development of mature, erythroid colonies from *Dot1L-*KO E10.5 YS cells was significantly impaired (Feng et al., 2010). However, the erythroid potential of the *Dot1L-*MM YS cells (based on numbers of colonies formed) appears to be relatively normal (**Figure 6A)**. To determine whether this difference in impairment was dependent DOT1L methyltransferase activity, *Dot1L-*MM E10.5 YS cells were plated in methylcellulose medium containing cytokines that promote *only* mature erythroid colony formation (mature BFU-E and CFU-E). Interestingly, no differences were found in colony number between *Dot1L-*MM and wildtype cells (**Figure 8I**). After 4 days in culture, mature erythroid cells were labeled with Annexin V and analyzed via flow cytometry for apoptosis. No differences were found in the percentage of cells undergoing apoptosis between mutant and wildtype cells (**Figure 8J**). These data are different than that of E10.5 *Dot1L-*KO YS cells. In contrast *Dot1L-*KO cells cultured in mature erythroid medium formed significantly fewer erythroid colonies compared to wildtype, and a greater percentage of these knockout erythroid cells were Annexin V-positive (Feng et al., 2010). In addition to the cell cycle analyses and apoptotic marker detection, Alkaline Comet assays were performed on day 6-8 WT and mutant YS ESREs to assess DNA damage.

Comet assays showed a greater DNA damage in *Dot1L-*MM compared to WT cells and the DNA damage was still higher in *Dot1L-*KO ESREs compared to MM (**Figure 8K**).

## 4. DISCUSSION

We previously demonstrated that DOT1L protein is essential for early hematopoiesis in mice (Feng et al., 2010). The gene trap mutation resulted in loss of the nucleosome binding domain as well as instability in mRNA level in *Dot1L-*KO mice (Feng et al., 2010). Loss of DOT1L protein led to lethal anemia in *Dot1L*-KO mice (Feng et al., 2010). In this study, we have shown that the histone methyltransferase activity of the DOT1L protein is also required for proper hematopoiesis during embryonic development. The enzymatic function of DOT1L, H3K79 methylation, proved essential for development of the myeloid, but ***not*** erythroid progenitors in the YS. Inactivation of the enzyme by a point mutation (Asn241Ala) in the catalytic domain of DOT1L protein results in YS progenitors’ failure to optimally undergo definitive hematopoiesis.

We observed that *Dot1L-*MM embryos had blood at E10.5, which suggests that the process of primitive hematopoiesis remained intact in the *Dot1L*-MM embryos. While the numbers of erythrocytes may be slightly reduced compared to wildtype embryos (**Figure 5C** and **6A**), the embryos clearly contained near-normal blood levels, and they were minimally anemic. This finding is quite different from the *Dot1l*-KO embryo.

Primitive erythroid cells begin to form in blood islands of the embryonic YS around E7.5, that provide oxygen support for the rapidly growing embryo (Frame et al., 2013). They continue to form in the YS until E9.0 (Baron et al., 2012) and enter circulation after the heart begins beating at E8.25 (Palis, 2016). Primitive erythropoiesis is vital for early growth and survival of the embryo, as embryos unable to carry out this process die by E9.5 (McGrath et al., 2015). Shortly after primitive erythroid cells form in the YS, a second wave of hematopoiesis begins in the YS, around E8.25 (Frame et al., 2013; McGrath et al., 2015). These second-wave progenitors are termed erythro-myeloid progenitors (EMPs) (Frame et al., 2013). EMPs begin seeding the fetal liver around E10.5-E11.5 (Baron et al., 2012; Frame et al., 2013; McGrath et al., 2015), where they attach to macrophages in erythroblastic islands and differentiate (Palis, 2008a). A third wave of hematopoiesis, which culminates in the production of definitive HSCs, occurs after YS EMP formation. HSCs are thought to originate primarily in the AGM region of the embryo but also possibly in the major arteries and placenta, from E10.5-11.5 (Baron et al., 2012; Frame et al., 2013). Both *Dot1L-*KO and *Dot1L-*MM, embryos die around E12.5, which suggests that the *Dot1L* methyl mutation has a minimal effect on the development of primitive erythroid cells. In contrast, the effect on the development of EMPs and HSCs are likely to be much more severe. Essential requirement of DOT1L for HSCs has also been demonstrated in adult hematopoiesis (Bernt et al., 2011; Jo et al., 2011b; Nguyen et al., 2011). Conditional *Dot1L* knockout (*Dot1L-*cKO) adult mice displayed defective hematopoiesis including bone marrow hypocellularity with decreased numbers of common myeloid (CMP), granulocyte/monocyte (GMP), megakaryocyte/erythrocyte (MEP), and HSC progenitors, pancytopenia, and eventual death (Jo et al., 2011b; Nguyen et al., 2011).

Hematopoietic differentiation assays were performed on E10.5 YS to assess how well *Dot1L-*MM mice could form definitive erythroid, myeloid, and multipotent, mixed progenitors. Although the mutants displayed a slight impairment in erythropoiesis and myelopoiesis, what was most striking was the near absence of mixed progenitor colonies in YS cultures (**Figure 6A**). This was also recapitulated in mESC differentiation assays (**Figure 3**). EMPs form in the YS beginning around E8.25 and are capable of forming multiple progenitors including macrophages, mast cells, basophils, neutrophils, and definitive erythroid progenitors (McGrath et al., 2015; Palis, 2016). This progenitors are early blood stem cells that arise before HSCs, and they leave the YS to colonize the fetal liver, where they proliferate exponentially and differentiate into definitive erythroid and myeloid progenitors (Palis, 2008b). The mixed colonies in the methylcellulose assays represent a good assessment of their presence and ability to proliferate in the murine YS. Given the lack of mixed colonies in the mutant YS, there clearly is an impairment in either EMP formation or the ability of the EMPs to proliferate and /or survive. If EMPs were not forming or surviving in the YS, then it is not surprising that there are no definitive erythroid or myeloid colonies present, since both definitive erythroid and granulocyte/macrophage colonies in the YS originate from EMPs. However, since definitive erythroid colonies grew from mutant YS cells in similar numbers as wildtype, and the fact that myeloid colonies do still grow (albeit at reduced numbers), EMPs are forming in the *Dot1L*-MM YS. In the mutant, it appears that there is an initial burst of EMP formation, but these stem cells then quickly differentiate into definitive erythroid and myeloid progenitors. Therefore, we hypothesize that there is impairment in the ability of *Dot1L* mutant EMPs to self-renew.

Interestingly, in inducible *Dot1L* knockout mouse models, *Dot1L*-deficiency leads to a particularly large decrease in the numbers of HSCs (Jo et al., 2011a). Other lineage-specific progenitors (common lymphoid, megakaryocyte/erythroid, granulocyte/macrophage, and common myeloid) are also significantly decreased (Jo et al., 2011a; Nguyen et al., 2011). In conditional mouse models of *Dot1L* deletion within the hematopoietic and endothelial compartments only, the earlier, multipotential hematopoietic compartments were more prominently affected (Bernt et al., 2011). *Dot1L*-mutants were able to form myeloid progenitors nearly as well as wildtype, but bipotential granulocyte/macrophage, LSK HSCs, and common myeloid progenitors were significantly decreased (Bernt et al., 2011). Megakaryocyte/erythroid progenitors were not significantly decreased (Bernt et al., 2011).

Additional support for this hypothesis is shown in the phenotype of the embryos themselves. E10.5 methyl mutant embryos contain blood and are only slightly anemic than wildtype embryos, but they still die by E13.5 (the majority by E12.5). Since EMPs colonize the fetal liver between E10.5 and E11.5 (McGrath et al., 2015) and proliferate exponentially there, the ability of EMPs to self-renew is necessary at this stage of development. It is conceivable that *Dot1L-*MM mice are dying by E12.5-13.5 because their EMPs cannot self-renew in sufficient numbers to seed the fetal liver. Studies in MLL leukemia cell lines add support to this apparent phenotype of rapid differentiation versus self-renewal in the absence of DOT1L-mediated H3K79 methyltransferase activity. When *Dot1L* was deleted from MLL-AF9 leukemia cells growing in methylcellulose culture, colonies were fewer in number, smaller, and displayed terminal differentiation in morphology, compared to the blast-like morphology seen in wildtype (Bernt et al., 2011). Likewise, similar changes were seen in DNMT3A-mutant AML cell lines. OCI AML2- and OCI AML3-DNMT3A-mutant cells treated with the DOT1L inhibitors SYC-522 and EPZ004777 seemed to undergo differentiation, as they had increased CD14 expression, a mature monocyte marker (Rau et al., 2016). Also, EPZ004777-treated OCI AML2 and OCI AML3 cells underwent RNA-seq analysis. Pathway analysis showed increased expression of genes involved in cell-cycle arrest, cell death, and differentiation (Rau et al., 2016).

The mechanisms dictating the lack of self-renewal are still being investigated. As DOT1L is known for involvement in transcriptional elongation (Nguyen and Zhang, 2011), it could be that transcription factors necessary for self-renewal are not being activated. Possible factors include LIF, interleukin-6, Sox17, or BMP4, all of which play roles in HSC self-renewal and expansion (Martinez-Agosto et al., 2007; Medvinsky and Dzierzak, 1998). Previous studies found that *Dot1L* knockout E10.5 KIT-positive YS progenitors had decreased GATA2 expression (Feng et al., 2010). GATA2 is important for definitive erythroid progenitor (such as HSC) proliferation and survival in response to growth factors (Kim and Bresnick, 2007; Medvinsky and Dzierzak, 1998). Thus, this decrease in GATA2 expression could prevent YS EMPs from self-renewing. It is also possible that these EMPs are more prone to cellular damage, such as DNA damage, cannot repair their genomes effectively, and differentiate as a result. Interestingly, several recent studies have documented a role for the DNA damage response in inducing differentiation of leukemic cells (Santos et al., 2014; Sherman et al., 2011). It is likely that accumulated DNA damage in mutant EMPs induces them to differentiate. *Dot1L-*MM YS cells plated in hematopoietic differentiation medium displayed G_0_/G_1_ cell cycle arrest (**Figure 8C-G**) and increased apoptosis compared to wildtype (**Figure 8 H**). Therefore, these progenitor cells are neither growing nor surviving. In addition to increased apoptosis and G_0_/G_1_ cell cycle arrest, mutant ESREs also displayed increased DNA damage in comet assays further suggesting that DOT1L methyltransferase activity plays a role in genome stability of early hematopoietic progenitor cells. Defective gene expression and DNA damage repair abnormalities could both be at play in the *Dot1L-*MM mice and cause the observed hematopoietic phenotype.

Perhaps the most striking result of these analyses is the apparent differences between the *Dot1L*-KO and the *Dot1L*-MM. Since the only defined activity associated with DOT1L is its role as a histone H3 methyltransferase, our data are consistent with the idea that there may exist methyltransferase-independent pathways for influencing differentiation in hematopoiesis, particularly erythropoiesis. The implications of this are far-ranging. First, this suggests that DOT1L possesses activities that are separable. If its tumorigenic activity can be isolated and safely targeted, leaving its essential regulatory role(s) intact, this would potentially have clinical implications. Second, these data suggest the possibility that there may be other functional domains of DOT1L protein that potentially are involved in the many diverse functions of this protein.

## Acknowledgements

This work was supported by the National Institutes of Health (grant R01DK091277). The mouse model was generated in the Transgenic and Gene Targeting Institutional Facility of the University of Kansas Medical Center, supported in part by the Center of Biomedical Research Excellence (COBRE) Program Project in Molecular Regulation of Cell Development and Differentiation (NIH P30 GM122731) and the University of Kansas Cancer Center (NIH P30 CA168524).

## Disclosure

The authors do not have any conflicts of interest.

## Notes

### Competing Interest Statement

The authors have declared no competing interest.

